# New mitochondrial genomes of leptosporangiate ferns allow modeling the mitogenomic inflation syndrome across all land plant lineages

**DOI:** 10.1101/2022.12.23.521604

**Authors:** Yanlei Feng, Susann Wicke

**Author notes:** Author for Correspondence: S. Wicke.

## Abstract

Plants’ mitochondrial genomes (mitogenome) evolve in a hard-to-predict fashion. To reconstruct the evolutionary trajectories of land plant mitogenomes, we, here, filled the last major mitogenomic gap within land plants by assembling the mitogenomes of the leptosporangiate ferns *Azolla filiculoides* and *Pteridium revolutum* and, secondly, analyzed the mitogenomic evolutionary regime shifts across land plants. By testing various Ornstein-Uhlenbeck stabilizing selection models in an a priori-free analysis of five selected mitogenomic traits, we observed 71 evolutionary regime shifts across 218 land plant species. These shifts can lead to genomic convergence, in which certain traits such as size, GC content, and the proportion of non-coding DNA converge, or non-converging regimes, which are characterized by exceptional paths of genomic evolution such as extreme GC content or size. We also found that non-seed plants have a slightly, but significantly, higher rate of synonymous substitutions across all gene classes than seed plants, and that ferns differ significantly in the number of nonsynonymous and synonymous changes compared with other non-seed and seed plants. This pattern matches an outstandingly high rate of RNA editing in the small but repeat-rich mitogenomes of leptosporangiate ferns. In sum, our study highlights the considerable changes in mito-chromosomal architecture that occur during land plant evolution and suggests that these changes may be related to increases in error-prone repair mechanisms. Further study of underrepresented plant groups such as ferns and lycophytes is needed to understand the mechanisms and dominating forces behind the evolutionary dynamics and the mitogenomic inflation syndrome.

**Significance Statement:** Our study provides new insights into the complexity and diversity of land plant mitogenome evolution and reveals that they take many turns of molecular evolutionary directions across 218 land plant species. Our results have the potential to inform future research in this area and to advance our understanding of the mitogenomic inflation syndrome during plant evolution.

## Introduction

Mitochondria are the powerhouse of eukaryotic cells. Their prime role is oxidative phosphorylation (OXPHOS) at the mitochondrial inner membrane, where chemical energy is generated through concerted action of five functional complexes. Complex I (NADH dehydrogenase, *nad* genes) and complex II (succinate dehydrogenase, *sdh* genes) harvest electrons from NADH and succinate, respectively. These electrons are then passed on to ubiquinone, complex III (cytochrome *bc*_*1*_ complex, *cob* gene), cytochrome c, whose biogenesis is mitogenome encoded in plants (*ccm* genes), and complex IV (cytochrome *c* oxidase, *cox* genes), resulting, eventually, in the reduction of oxygen to water. An associated mitochondrial ATP synthase complex (*atp* genes) uses the energy released from the redox reactions to form ATP.

Following their endosymbiotic uptake, most genes of the proteobacterium-like prokaryote (Gray *et al*., 1999; Martijn *et al*., 2018) have migrated to the nucleus, leaving only a functionally reduced, now semi-autonomous genome in the organelle. Unlike in metazoans and fungi, whose gene sets are conserved and length variations originate from differences in tRNA and intron content or lengths (Gissi *et al*., 2008; Sandor *et al*., 2018), mitogenome evolution in the green lineage is particularly difficult to predict. For example, in green algae, there is a shift from circular to linear genome structures and an extensive size variation (Vahrenholz *et al*., 1993; D. R. Smith *et al*., 2010; David Roy Smith *et al*., 2010; Lee and Hua, 2014; Zheng *et al*., 2018). Some chlorophyte algae contain only seven protein-coding genes (Burger and Nedelcu, 2012; Turmel *et al*., 2013), whereas charophyte algae, the sister group of land plants possess gene-rich mitochondrial genomes (Turmel *et al*., 2013). With between 66 kb and 11 Mbp in size and partly split into multimeric chromosomes, land plants exhibit the extremes of mitogenome evolution. Bryophyte mitogenomes appear in a state of structural stasis (Liu *et al*., 2014), whereas “higher” plant mitogenomes are often size-inflated with an unpredictable genomic configuration (Knoop *et al*., 2011; Mower *et al*., 2012). Higher plant mitogenomes also tend to integrate DNAs originating either from other genome compartments within the cell or horizontally from another species. The capricious dynamics of plant mitogenomes hamper their study. So, complete mitogenomic sequences are still unavailable for leptosporangiate (true) ferns. Hence, it remains unclear to date when and why mitogenomic evolutionary patterns change during the evolution of land plants.

This work aims to improve our understanding of the course and direction of mitogenomic reconfigurations during land plant evolution and to provide a resource for future plant mitogenome studies. In so doing, we close an important taxonomic gap by presenting the first fully assembled and annotated mitochondrial genomes of two leptosporangiate (true) ferns. Using phylogenomic comparative analyses, we analyze patterns of gene losses, DNA incorporations, structural dynamics, and molecular rates of evolution across land plants. The results of our work show that leptosporangiate ferns are among the highest ranking of extraordinary mitogenomes. We hypothesize that the independent departures from evolutionary stasis in vascular plants underlie a common cause that dates back to the transition from a dominant gametophytic to a dominant sporophytic phase.

## Materials & Methods

### Data acquisition and processing

We accessed 819 entries from 309 species/subspecies mitogenomes of land plants and charophyte algae available in the public genome repository of NCBI GenBank. As these data included draft mitogenomes, we filtered the dataset and included only some but not all of these for our analyses (Table S1). Since the annotations of the genomes were not uniform, we corrected gene names from “*orf*” denominations to the accepted ones (e.g., *orfB* – *atp8, orf25* – *atp4, orfX* – *mttB*). In several cases, we noticed that exons had been missed or intron/exon boundaries were incorrect in the original annotations of the multi-intronic *cox2, nad1, nad2, nad4, nad5*, and *nad7*. We corrected these errors manually using GENEIOUS v11 (https://www.geneious.com) after inspecting all original annotations. We also marked cases of potential mis-assemblies or incomplete assemblies (Table S1). To obtain a first overview of the land plant mitogenome landscape, we plotted scattergrams among groups using the GGPLOT2 package (Gómez-Rubio, 2017).

### Assembly and annotation of new mitogenomes

We assembled and annotated the complete mitogenomes of two leptosporangiate ferns by reanalyzing paired-end genome-shotgun and transcriptome data of the heterosporous water fern *Azolla filiculoides* (Brouwer *et al*., 2017; Dijkhuizen *et al*., 2018; SRA accession numbers: ERR2503876–ERR2503878, ERR2114811) and the terrestrial fern *Pteridium revolutum* (Liu *et al*., 2019; SRA accessions: SRR7121815; SRR6920707). Transcriptome data were used to assist the annotation procedure and to analyze RNA editing. Similarly, we reanalyzed the mitogenomes of *Cuscuta australis* (Zhuang *et al*., 2018; Sun *et al*., 2018; SRA accessions: SRR5851367, SRR5934903, SRR7719072, SRR7719078), *Cuscuta campestris* (Vogel *et al*., 2018; SRA accessions: ERR1913332, SRR8669724, SRR8669805, ERR1916364), and *Ipomoea nil* (Wei *et al*., 2015; Hoshino *et al*., 2016; SRA accessions: DRR024544, DRR024546, SRR1772255) to compare RNA editing patterns.

We cleaned all raw DNA and RNA data of low quality bases using TRIMMOMATIC v0.36 (Bolger *et al*., 2014). *De novo* assemblies of every DNA data set were performed with SPADES v3.13 (Bankevich *et al*., 2012). By using a custom-compiled reference set of conserved mitochondrial genes, we extracted all contigs of a potential mitochondrial origin with MEGABLAST (Camacho *et al*., 2009) searches, with default settings. To extend the contig flanks, we iteratively backmapped the cleaned paired-end reads and connected overlapping contigs until no more connection anchors were found. Even coverage and the correct assembly of the mitogenome drafts were re-confirmed by read-mappings and inspecting potential read pair splits by eye.

For the annotations of the newly assembled mitochondrial sequences, we predicted the gene models by comparisons with published relatives. For the fern mitogenome, annotation was assisted by RNASeq data, from which also RNA editing sites were assessed. For the latter, we mapped cleaned RNASeq reads to the most likely gene models in GENEIOUS v11 (settings: “medium sensitivity”). We traced polymorphisms between DNA and RNA data using the SNP variant plugin of GENEIOUS and a minimum per-site coverage of one and a minimum variant frequency of 25 %. The herein used RNASeq data was not optimized originally for capturing organelle transcripts. Nevertheless, mitogene transcripts were evenly present. Discrepancies between DNA and RNA data and the potential RNA editing sites were inspected by eye, sorted, and summarized using a custom Perl script (Mendeley Data, doi:10.17632/4fgcswrw3v.1).

### Plastid-derived and dispersed repeat sequences

Plastid-derived DNA in mitochondrial genomes (hereafter: MTPT) was determined through MEGABLAST (evalue filter: 1E^-5^) against a custom plastid genome (plastome) database, consisting of plastid data from each of the species studied here; if a species-specific plastome was unavailable, we replaced it with that of a close relative (Supporting Table S1). BLAST hits of less than 100 bp were masked. We also used self-self MEGABLAST searches to locate dispersed mitochondrial repeats. Hits with an identity below 95 % and a length shorter than 100 bp were ignored. The number and length of MTPT and repeats per species were evaluated by a custom Perl script (Mendeley Data, doi:10.17632/4fgcswrw3v.1). Overlapping repeat motifs were counted only once. To estimate the dispersed repeat length, we considered only the length of the duplicated sequences.

### Analysis of molecular evolution

We aligned all coding regions of charophytes and land plants using a translation alignment in combination with MAFFT v7.036b (Katoh and Standley, 2013) with automated selection of the best alignment options. Duplicated and foreign genes were excluded. Where data from more than one accession per species were available, we retained only one mitogenome record for the analysis of molecular evolution. Also, to improve the balance of the lineage representation in our data, we limited the lineage representation to maximum four species of the same genus (Table S1). One aim of this work was to analyze changes on the DNA level and match these to the extent of RNA editing. Therefore, we excluded no sites from gene data sets, even though RNA editing is a given.

To resolve unclear phylogenetic placements of some lineages or single taxa for subsequent (molecular) evolutionary modeling, we computed a phylogenetic tree from the concatenated codon dataset of all mitochondrial genes using RAxML (Stamatakis, 2014) under the GTR-GAMMA model. Node support for the best tree was inferred from 1 000 bootstrap replicates using the same model. The resulting phylogenetic tree was the basis for all subsequent tree-based analyses.

We modeled the structural evolution of plant mitogenomes using Ornstein-Uhlenbeck stabilizing selection models (Ingram and Mahler, 2013). This *a priori*-free approach uses stepwise Akaike Information Criterion to infer evolutionary regimes of trait evolution across phylogenies. To do so, this method locates shifts of mitogenomic trait regimes (mitogenome size, GC content, length of coding regions as proxy for gene content changes, proportion of non-coding DNA, and repetitive DNA) in our land plant phylogeny by stepwise addition until the model fit improves no further. Next, it collapses the best regimes to identify convergences in the data. For computational reasons, we allowed a maximum of 80 primary regimes across our 218 taxa tree, and inferred the convergence regimes from all instead of only the best ones.

For every mitochondrial protein-coding gene, we used the HYPHY suite v2 (Pond et al., 2005) to compute the rates of nonsynonymous and synonymous substitution (dN and dS, respectively) as described in detail earlier (Wicke *et al*., 2014; Wicke *et al*., 2016). In brief, using the charophyte *Klebsormidium* as an outgroup, we estimated dN and dS for every mitochondrial gene by pairwise relative rate analysis using the MG94 codon model under the maximum likelihood paradigm and likelihood ratio testing. We visualized differences in molecular evolutionary rates with biplots, for which dN and dS were averaged per species and gene (Wicke and Schneeweiss, 2015). Bagplots were employed to illustrate lineage-specific molecular evolutionary rates, complemented by unpaired Wilcoxon testing with sequential alpha error correction to assess the significance of group-specific rate variation (detailed in Wicke *et al*., 2014). For an alternative visualization of mitochondrial rate evolution, we also fitted and optimized an unconstrained likelihood function (LF) that uses a full MG94 model (with local parameters for both dN and dS) to the specified phylogeny of land plants (Supporting Figure S1) and the combined (concatenated) mitochondrial gene data.

Associations between mitochondrial rates of evolution, genomic traits such as mitogenome-wide GC content, genome size, the ratio of coding to non-coding DNA, repeat density (Table S1), and processes underlying the nucleotide substitution process (transition: transversion ratio, GC content of coding sequences, which were co-estimated from the data during analysis) were evaluated in a Bayesian framework with COEVOL v1.4 (Lartillot and Poujol, 2011). We used a fixed land plant topology and a concatenated data set of mitochondrial coding sequences to set up two parallel analyses to estimate (co-)correlations between dS and the various genomic traits. We ran COEVOL with one chain each, from which every tenth generation was kept until 30 000 effective samples were collected. MCMC chain convergence was checked by computing the discrepancies and effective sample sizes of the posterior averages, tree lengths, and several other summary statistics obtained from the independent runs. Per run, the posterior averages were computed after removing the initial 10 % of samples per chain as burn-in.

## Results and Discussion

Based on data from seven charophyte algae (as outgroup), 15 liverworts, 46 mosses, four hornworts, three lycophytes, two eusporangiate and two leptosporangiate ferns, as well as 241 seed plants (Supporting Material: Table S1), we (re-)analyzed and modeled the trajectories of mitogenome evolution (Figure 1). To do so, we first reconstructed the phylogenetic relationships of our study taxa as basis for all subsequent phylogeny-aware statistical analyses. This phylogenetic reconstruction placed hornworts as sister to liverworts and mosses in a monophyletic bryophyte clade (Figure 1; node support in Figure S1) as the earliest-branching lineage of land plants. These findings are in line with other recent phylogenetic hypotheses of land plant evolution based on transcriptome wide-inferences (e.g., Wickett *et al*., 2014; One Thousand Plant Transcriptomes Initiative, 2019; Nishiyama *et al*., 2018; Li *et al*., 2020).

**Figure 1.**
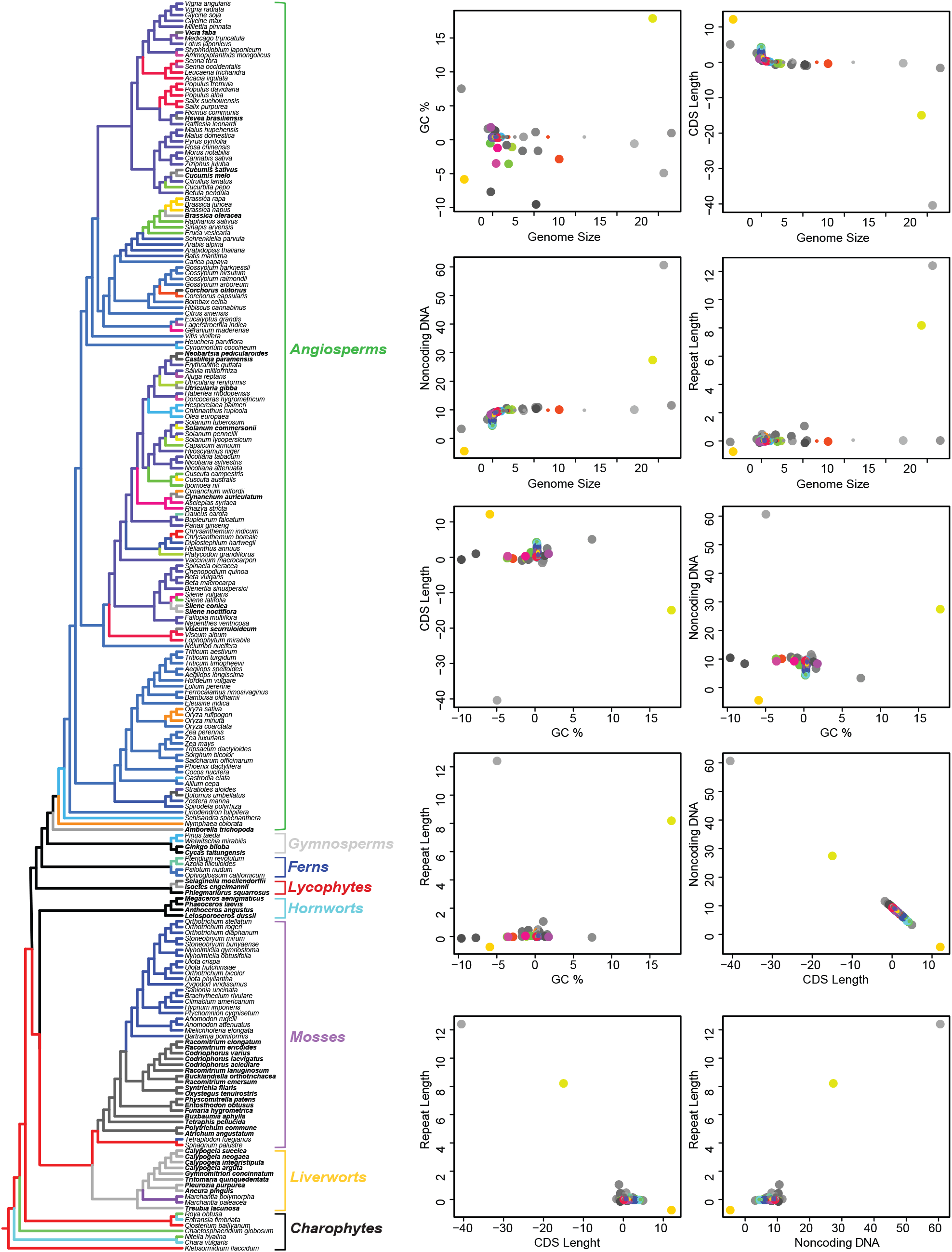
Mitogenomic evolutionary regimes in land plants. The phylogenetic tree of land plants on the lefthand side highlights the distribution of converging and non-converging mitogenomic evolutionary regimes. Converging regimes are indicated by same colors across the various branches; non-converging regimes are illustrated in gray tones of branches and boldface taxa labels. Point plots to the right show the relative associations of the various converging and non-converging regimes between all trait pairs.

### Evolution of mitogenome structure

We tested 434 Ornstein-Uhlenbeck stabilizing selection models in an *a priori*-free analysis of major mitogenomic traits, allowing us to identify 71 genome-evolutionary shifts and 40 prime regimes of structural mitogenomic evolution in land plants (Figure 1). Of these, 53 shifts were directed towards one of 22 convergence regimes, in which the set of the five analyzed structural traits are highly similar. For examples, the mitogenomic structure regime, defined here by a combination of mitogenome size, nucleotide composition, the length of protein-coding genes, the amount of repetitive DNA, and the proportion of non-coding DNA, that characterizes the eusporangiate ferns *Ophioglossum* and *Psilotum* is seen also in many angiosperms, including most monocots and several eudicots (Figure 1; Supporting Table 2). The newly assembled leptosporangiate ferns *Azolla* and *Pteridium*, however, belong to a much rarer regime of mitogenomic evolution, shared only with a single eudicot species (*Daucus carota*, Apiaceae). At regime shifts, the rate to evolve towards another optimum differs largely between traits (Table 1). Nucleotide composition adapts two orders of magnitudes faster to the trait optimum than mitogenome size and the proportions of protein-coding, non-coding, and repetitive DNA. However, high rates of stochastic trait evolution of the (correlated) lengths of the mitogenome and its repeated DNA proportion reduce the distinctness of the regimes (σ^2^ = 9.31E+09 and 1.07e+06, respectively, compared with σ^2^ = 2.21e+04, 3.27e+04, and 9.91E-02 for protein-coding and non-coding DNA lengths and GC content).

**Table 1.** Variance of mitogenomic traits in land plants.

| Phylogenetic ANOVA (n = 218 taxa) |  |  |  |  | Posthoc test of group mean differences |  |
| --- | --- | --- | --- | --- | --- | --- |
| Response | Sum Sq | Mean Sq | F value | Pr(>F) | Pairwise t-values | Pairwise corrected P-values |
| *Bryophytes vs. vascular plants* |  |  |  |  |  |  |
| Mitogenome length | 1206.745 | 1206.7453 | 14.2769 | 0.767 |  |  |
| % GC | 0.055043 | 0.055043 | 94.892498 | 0.453 |  |  |
| % repetitive DNA | 2281.157 | 2281.15692 | 33.79413 | 0.663 |  |  |
| % noncoding DNA | 22355.212 | 22355.2119 | 777.141006 | 0.014 | 27.87725 | 0.014 |
| % protein-coding DNA | 15038.266 | 15038.266 | 753.91865 | 0.018 | -27.45758 | 0.018 |
| *Non-seed vs. seed plants* |  |  |  |  |  |  |
| Mitogenome length | 1280.057 | 1280.05735 | 15.207359 | 0.779 |  |  |
| % GC | 0.022665 | 0.022665 | 30.838068 | 0.684 |  |  |
| % repetitive DNA | 937.8214 | 937.82142 | 12.685431 | 0.787 |  |  |
| % noncoding DNA | 20166.32 | 20166.3195 | 513.933843 | 0.115 |  |  |
| % protein-coding DNA | 13437.01 | 13437.0102 | 486.701612 | 0.095 |  |  |

**Table 2.** Coevolutionary associations between mitogenomic traits based on 30 000 effective samples.

|  | dS | Ts/Tv | GC | Size | % protein-coding | % struct. RNA | % non-coding | % repeat length |
| --- | --- | --- | --- | --- | --- | --- | --- | --- |
|  | Fully controlled correlations |  |  |  |  |  |  |  |
| **dS** | - | 0.66 | 0.95 | 0.18 | 0.23 | 0.01 | 0.30 | 0.19 |
| **Ts/Tv** | 0.50 | - | 0.01 | 0.55 | 0.73 | 0.59 | 0.20 | 0.87 |
| **GC content** | 0.87 | 0.01 | - | 0.11 | 0.08 | 0.84 | 0.54 | 0.81 |
| **size** | 0.01 | 0.03 | 0.95 | - | 0.01 | 0.90 | 0.01 | 1.00 |
| **% protein-coding** | 0.96 | 0.97 | 0.03 | 0.01 | - | 1.00 | 0.01 | 0.10 |
| **% struct. RNA** | 0.80 | 0.98 | 0.05 | 0.01 | 1.00 | - | 1.00 | 0.57 |
| **% noncoding** | 0.01 | 0.04 | 0.97 | 1.00 | 0.01 | 0.01 | - | 0.09 |
| **% repeat length** | 0.06 | 0.84 | 0.94 | 0.99 | 0.01 | 0.01 | 0.99 | - |
|  | Marginal correlations |  |  |  |  |  |  |  |

We identified 18 non-converging regimes of structural mitogenomic evolution. Those regimes express exceptional paths of genomic evolution, e.g., due to extremes in the GC content, size, or the lengths of protein-coding or non-coding DNA (Figure 1). Only a handful of flowering plants experienced unmatched evolutionary trajectories of mitogenomic evolution. Those plants include the basal angiosperm *Amborella* (Rice *et al*., 2013), one in 28 monocots, as well as six of 61 rosid representatives and seven of 70 asterid species. Of the latter, three species are hemiparasitic (*Viscum scurruloideum* [Viscaceae], *Neobartsia pedicularis, Castilleja paramensis* [both Orobanchaceae]), and one is a carnivorous plant (*Utricularia gibba*, Lentibulariaceae). In contrast, none of the five (physiological) holoparasites stick out in terms of mitogenomic structures, including the *Cuscuta* species (Supporting Figure S2 and Table S1).

It is obvious that mitogenomic evolution experienced many independent shifts across and within the various land plant lineages (Figure 1). The distribution of regimes shows that reconfiguration of mitogenome structure seems to have occurred more ancestrally within bryophytes than in flowering plants. Evolutionary patterns remain unclear in early vascular plants and gymnosperms, likely due to taxonomic underrepresentation. Group-wise comparisons using phylogenetic ANOVA show that the proportion of non-coding and the length of protein-coding DNA differs significantly between bryophyte lineages and vascular plants (phylAnova P-values: 0.014 and 0.018, respectively; n=211). Further hypothesis testing using phylogenetic comparative methods under the maximum likelihood paradigm corroborate that non-coding and protein-coding DNA of non-vascular plants differ notably from vascular plants in their architectural traits (Likelihood ratio test (LRT): 2Δ = 1 699 and 623, df = 1, both P-values < 0.001). Both traits apparently have evolved gradually rather than punctually (LRT: 2Δ = 509 and 674, df = 1, both P-values < 0.001). In line with this finding, time (long paths) contribute more than short paths to the emergence of extreme trait values (LRT: 2Δ = 588 and 1 313, df = 1, both P-values < 0.001). In contrast, no statistically significant groupwise differences in GC content or the amount of repetitive DNA between bryophytes and seed plants exists. We also detected no group-specific differences between non-seed and seed plants in any of the analyzed mitogenomic traits (Table 1). Together, these results suggest that a primary mitogenomic disturbance might have occurred along the evolution of vascular plants that manifested over time as genomic drift rather than adaptive change.

### Lineage-specific changes of mitogenomic architecture

Bryophyte mitogenomes are normally circularized molecules of 143 to 187 kilobases (kb) in liverworts and 100 to 141 kb in mosses (Supporting Table S1). Compared to charophytes, the GC content of mosses and liverworts are slightly elevated (Figure 2) and the mito-chromosomal synteny in both these groups is highly conserved (Liu *et al*., 2014; but see Myszczynski *et al*., 2018). Hornworts have larger mitogenomes (185–242 kb) than mosses and liverworts, and they show notable inversions or translocations (Wang *et al*., 2018). We observe hardly any repeats in bryophyte mitogenomes, with mosses being particularly poor in repeats longer than 100 bp (Figure 2; Table S1).

**Figure 2.**
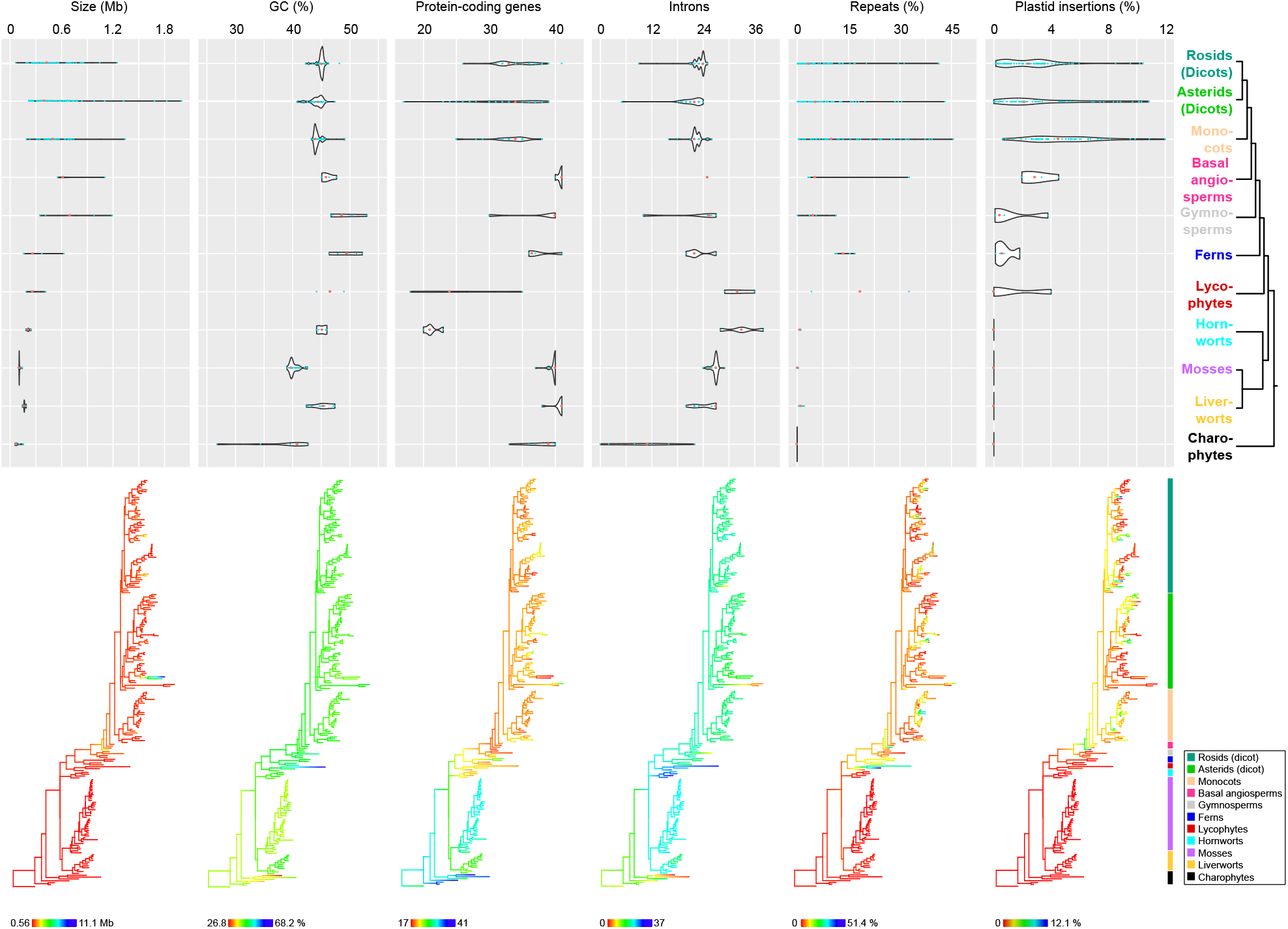
Mitogenomic trait changes across land plants. Changes of mitogenomic size, GC content and coding capacity are visualized by lineage using violin plots, which emphasize the trait-specific medians per lineages, whereas the macroevolutionary shifts are graphically summarized over the land plant tree. The trait range with minimum and maximum are indicated below the tree; for simplicity, taxa names are omitted. A lineage tree or color bar to the righthand side of both the violin plots and trait trees, respectively, indicate the position of the various land plant lineages.

In vascular plants, mitogenome size increases gradually. Alongside the genomic expansion, repetitive and foreign DNAs accumulate (Figure 1 and 2) and multi-chromosomal configurations of mitogenomes emerge (Table S1). The number of mitochondrial chromosomes or sub-molecules (“episomes”) can be in the hundreds (Sloan, Alverson, Chuckalovcak, *et al*., 2012). As a result, mitogenomic synteny breaks, leaving only a few collinear gene clusters intact (Richardson *et al*., 2013). Despite a trend towards larger genomes in lycophytes (*Isoëtes*: 187 kb, *Selaginella*: 261 kb, *Huperzia*: 414 kb), these early-diverging vascular plants show a considerable variation in gene content and the proportion of noncoding DNA (Supporting Table S1). Coinciding with structural rearrangements, nucleotide compositional biases also become extreme in lycophytes, with the spikemoss *Selaginella* reaching an outstanding 68% (Hecht *et al*., 2011).

Mitogenome expansion continues in eusporangiate ferns (*Ophioglossum*: 372 kb, *Psilotum*: 629 kb). In contrast, a considerable size reduction characterizes the two leptosporangiate ferns *Azolla* and *Pteridium* (both ∼157 kb; Supporting Figure S2), which still are rich in dispersed repeats. The increasing proportion of repetitive DNA is especially obvious in comparison with bryophytes, lycophytes, and eusporangiate ferns.

In seed plants, mitogenome structure varies most dramatically (Figure 1). Mitogenome size inflates (median lengths: 400 kb; Figure 2) up 11 Mbp in the eudicot *Silene* and Gymnosperm Siberian larch (Mower *et al*., 2012; Sloan, Alverson, Chuckalovcak, *et al*., 2012; Putintseva *et al*., 2020). However, in light of the vastly understudied ferns and gymnosperms, it remains unclear whether this flowering plant lineage represents a global or a local maximum of mitogenome evolution in land plants. The evolutionarily older cycads and *Ginkgo* all have smaller genomes than the investigated conifer representatives (Figures 1 and 2; Supporting Table S1). Also basal angiosperms are more conserved than the evolutionarily younger rosid and asterid superclades. Compared with the frequent expansions or contractions of dicot mitogenomes, monocot mitogenomes are rather conserved in size. A notable exception from the latter is the nonphotosynthetic orchid *Gastrodia* and *Cyperus esculentus* (Yuan *et al*., 2018; Niu *et al*., 2022), which represent rare monocot mitogenomes of over one million base pairs (Figure 1; Table S1).

An extreme physical reduction of the mitochondrial genome occurred in mistletoes. With 66 kb, *Viscum scurruloideum* retains the smallest land plant mitogenome (Skippington *et al*., 2015). European mistletoe (*V. album*) shows no such extreme mitogenomic downsizing (Petersen *et al*., 2015; Skippington *et al*., 2017) and also belongs to a rather common selective regime of mitogenome evolution (Figure 1). A similar phenomenon also occurred in Balanophoraceae, where the mitogenome of *Rhopalocnemis phalloides* is 131 kb in size, whereas that of *Lophophytum mirabile* is inflated to 822 kb with a remarkable extent of integrated foreign sequences (Schelkunov *et al*., 2019; Yu *et al*., 2022; Sanchez-Puerta *et al*., 2019). Considering other, independently evolved parasitic plant lineages *s*.*l*. (*Cynomorium*, Cynomoriaceae: Bellot *et al*., 2016; *Castilleja* and *Neobartsia*, Orobanchaceae: Fan *et al*., 2016; *Rafflesia*, Rafflesiaceae: Sanchez-Puerta *et al*., 2017; *Gastrodia*, Orchidaceae: Yuan *et al*., 2018; *Hypopitys monotropa*, Ericaceae: Shtratnikova *et al*., 2020; *Cuscuta* spp.: Anderson *et al*., 2021; Lin *et al*., 2022; *Aeginetia indica*, Orobanchaceae: Choi and Park, 2021), Hypopitys monotropa there is no clear trend regarding heterotrophy-associated mitogenome evolution. The nonphotosynthetic *Cynomorium, Gastrodia, Hypopitys*, and *Cuscuta japonica* have larger mitogenomes than their closest photosynthetic relatives. Other *Cuscuta*, as well as *Aeginetia* and two hemiparasites belonging to Orobanchaceae show mito-structural peculiarities similar to many other angiosperms (Table S1). It is worth noting that, besides some Viscaceae, also mitogenomic evolution in Orobanchaceae parasites show no convergence with that of other land plants.

### Changes of intron distribution

Two structurally different types of self-splicing and mobile introns reside in land plant mitochondrial genes (mitogenes): Group I introns that are spliced in a linear form and group II introns that require a lariat shape for splicing (Saldanha *et al*., 1993). Their evolution has been the focus of many studies, showing a dominance of group II introns (see Mower, 2020 for a detailed review). Liverworts, mosses, and hornworts show little within-lineage variation of intronization patterns (Figure 2 and Table S1). Between groups, intron types and numbers differ notably. Over 80 introns were identified in bryophytes, but none is conserved across the three lineages (Knoop, 2010; Knoop, 2013). Introns are conserved between the leptosporangiate ferns *Azolla* and *Pteridium*, but their intronization pattern is in stark contrast to that of other land plant lineages (Table S1). The two true fern species possess intronized genes for the large ribosomal RNA subunit (*rrnL*) (Supporting Figure S3), which differ between the two species and to the *rrnL* introns of liverworts and charophytes. RNAseq data indicates that all introns are apparently spliced efficiently and correctly (Supporting Figure S3). The single intron in *Azolla* corresponds to the first intron of *Pteridium*, most likely originating from the same homing event (Figure S3). *Pteridium* contains three more introns in *rrnL*, which leads to two extremely short *rrnL* exons (8 and 31 bp in length, respectively).

Perhaps as a consequence of an elevated recombination rate, splicing shifts from *cis* to *trans* can occur. In lycophytes (Grewe *et al*., 2009; Hecht *et al*., 2011) and gymnosperms (Guo *et al*., 2020), the shifts occurred more frequently and irregular among genes. In angiosperms, *trans*-splicing introns are mostly limited to *nad1, nad2*, and *nad5*, except for some shifts, such as in *ccmFc* of Convolvulaceae (Lin *et al*., 2022). Interestingly, so far there is no *trans*-splicing intron found in ferns, indicating that the *trans*-splicing introns in lycophytes and gymnosperms evolved independently.

### Changes in the proportion of foreign DNAs

Many mitogenomes contain fragments of nuclear or plastid origin. Although nuclear mitochondrial inserts can make up half the length of the mitogenome (*Arabidopsis*: Marienfeld *et al*., 1999; *Oryza*: Notsu *et al*., 2002; *Beta*: Satoh *et al*., 2006; Cucurbitaceae: Alverson *et al*., 2010 and ; Rodríguez-Moreno *et al*., 2011; *Malus*: Goremykin *et al*., 2012; Fabaceae: Alverson *et al*., 2011; bryophytes: Liu *et al*., 2014), the direction of transfer is often unclear. In contrast, plastid-derived DNAs (MTPT) are particularly frequent in mitogenomes (Figure 2) and easy to detect, because plastid genomes typically select against incorporations of non-native DNA (Wicke *et al*., 2011; but see Smith, 2014).

The length and number of MTPT differ largely between the various land plant lineages (Table 1). Although present in *Isoëtes*, lycophytes tend to have no detectable DNAs of plastid origin—similar to bryophytes (Hecht *et al*., 2011; Liu *et al*., 2012). In ferns, MTPT are common, albeit low in number and short in length (max. 3 kb; Figure 2 and Supporting Table S1). In seeds plants, including *Cycas*, plastid-derived DNAs are widespread and make up a considerable portion of mitogenome lengths (Chaw *et al*., 2008) and can exceed 10 kb (Stern and Lonsdale, 1982; Alverson *et al*., 2010) (Table S1). The transferred plastid DNAs are not region-specific, although some are apparently more likely to be incorporated than others (Wang *et al*., 2018).

Plant mitogenomes are also known to incorporate DNA from distantly related species (horizontal DNA transfer, HGT). Horizontally transferred DNAs are not known from bryophytes, but vascular plant mitogenomes seem to be particularly prone to HGT. Lycophytes and ferns contain some mitogenomic tRNA genes of bacterial origin (Guo *et al*., 2017), and the eusporangiate fern *Botrychium* has acquired DNAs horizontally from flowering plants (Davis *et al*., 2005). In contrast, we found no evidence for horizontally acquired fragments in the mitogenomes of *Azolla* and *Pteridium*. Especially parasitic plants are best known for their host-derived mitochondrial genes (Davis and Wurdack, 2004; Barkman *et al*., 2007; Sanchez-Puerta *et al*., 2017; Skippington *et al*., 2017; Sanchez-Puerta *et al*., 2019) and at least one case of parasite-to-host transfer (Mower *et al*., 2004). Similar to *Viscum*, some *Cuscuta* spp. are apparently devoid of horizontally acquired or host-derived DNA (Anderson et al., 2021; Lin et al., 2022). These data suggest that the extent of mitogenomic HGT follows highly lineage-specific routes.

### Evolution of mitochondrial coding capacity

The semi-autonomous genetic system of plant mitochondria works in concert with elements encoded in the nuclear genome. Land plants retain between 16 and 43 protein-coding genes in their mitogenomes (Figure 2; Table S1). Hardly any changes in the coding capacity for genes of the respiratory complex occur during land plant evolution (Figure 2 and 3). In contrast, losses of genes for mitochondrial translation occur multiples times independently (Figure 3). Land plant mitogenomes gain genes, too. The highly dynamic mitogenome structure offers different ways to create paralogs, novel genes, or re-gain a formerly lost gene (Wynn and Christensen, 2018). Also, intracellular gene transfer, horizontal gene transfer, gene conversion, retroprocessing, and *de novo* evolution of genes are potential mechanisms to yield “novel” or “chimeric” mitogenes.

**Figure 3.**
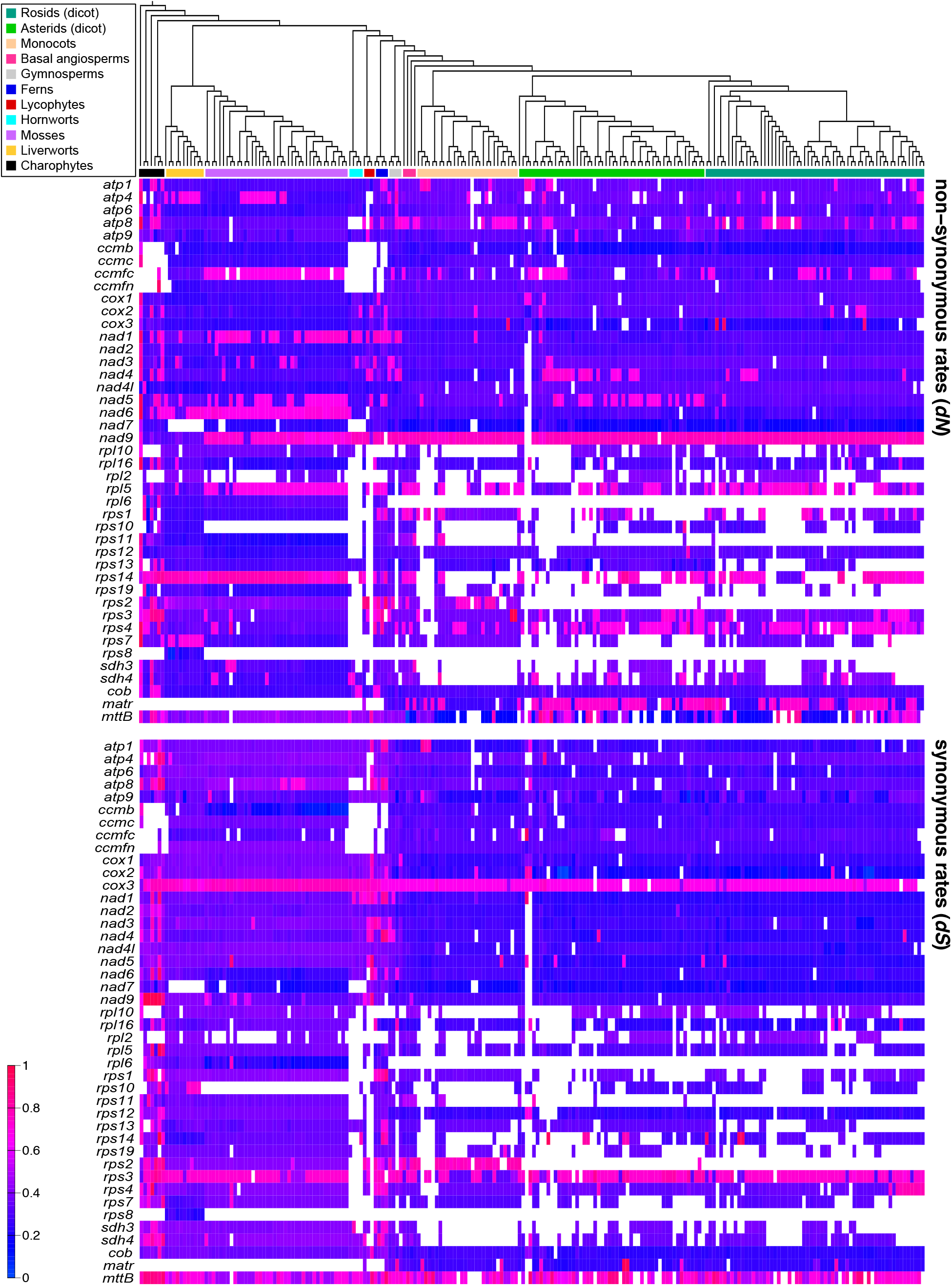
Nonsynonymous (dN) and synonymous rates (dS) of mitochondrial protein-coding genes in land plants. Graphic summary of relative dN (upper panel) and dS (lower panel), showing the ups and downs of molecular evolutionary rates for all mitogenes across 218 land plant taxa and charophytes. High molecular evolutionary rates are shown in red, whereas blue indicates low rates. Blank fields indicates gene losses or artificial errors caused by uncorrected misassemblies or misannotations.

#### Genes for oxidative phosphorylation

Plant mitogenomes encode up to 24 genes for the OXPHOS complexes: *nad1*-*7, 9*, and *4L, sdh3* and *sdh4, cob* for apocytochrome *b* plus *ccmB*/C/Fc/*Fn* for cytochrome *c* biogenesis, *cox1*-*3, atp1, 4, 6, 8*, and *9*. Among these OXPHOS-related genes, *sdh* genes are most frequently lost in monocots and eudicots (Figure S3 and Table S1). Although *sdh3* and *4* are normally conserved in non-seed plants and basal angiosperms, hornworts (Villarreal *et al*., 2018) and the enigmatic gnetophyte *Welwitschia* (Guo *et al*., 2016) retain only degraded copies of *sdh3*. Their frequent losses may be the results of functional gene transfer (Adams *et al*., 2001).

Losses of other OXPHOS-related mitochondrial genes are rare (Figure 3) and mostly explainable by prior functional transfer of a mitogene to the nucleus. *Atp8* is a pseudogene in hornworts (Villarreal *et al*., 2018) and *atp4* is missing from the lycophyte *Selaginella* (Hecht *et al*., 2011). The absence of *cox2* with no apparent functional consequences was reported for *Vignia* (Alverson *et al*., 2011; Naito *et al*., 2013). *Nad7* is physically or functionally lost several times independently within liverworts (Groth-Malonek *et al*., 2007; Liu *et al*., 2011), mosses (Bell *et al*., 2014; Liu *et al*., 2014; Goruynov *et al*., 2018), and hornworts (Li *et al*., 2009; Xue *et al*., 2010; Dong *et al*., 2018; Villarreal *et al*., 2018). The fern *Pteridium* seems to have lost *nad9*. We failed to retrieve contigs with similarity to full-length *nad9* orthologues in assembled whole-genome shotgun and transcriptome data. An additional cross-species read-mapping analysis using the *nad9* gene of *Azolla* as reference identified *Pteridium* reads for only the gene’s 5’-end. Based on low-coverage genomic sequencing (∼ 7 X), *nad4L* was recently reported as a pseudogene in the hemiparasitic angiosperm *Krameria* (Zervas *et al*., 2019). The most extreme losses of *nad* genes characterize some members of the Viscaceae family (Petersen *et al*., 2015; Skippington *et al*., 2015; Zervas *et al*., 2019)(Skippington *et al*., 2015; Petersen *et al*., 2015; Skippington *et al*., 2017; Zervas *et al*., 2019). All *nad* genes of *Viscum* and *Phoradendron liga*, another hemiparasitic mistletoe of Viscaceae, are either absent from their mitogenomes or highly divergent. Recent studies confirmed the lack of complex I proteins in European mistletoe (*V. album*); all other complexes were dramatically rearranged (Senkler *et al*., 2018; Maclean *et al*., 2018). *Viscum* also lacks an intact *ccmB* gene (Petersen *et al*., 2015). The *ccmB* copy residing in the *V. album* mitogenome might have been acquired from another species (Skippington *et al*., 2017).

With the evolution of charophyte algae and land plants, the *F* subunit of the cytochrome *c* biogenesis complex, encoded by a single gene in the charophyte *Klebsormidium* (Valach *et al*., 2014), breaks into two (*ccmFc* and *Fn*). During the evolution of land plants, *ccm* genes are lost multiple times and often in concert. The liverwort *Treubia lacunosa* retains *ccmB* and *ccmFc* as pseudogenes, while subunits *c* and *Fn* are physically lost (Liu *et al*., 2011). Notably in Brassicaceae, *Allium* (Poaceae), and *Trifolium* (Fabaceae), *ccmFn* is split into two separated, apparently not *trans*-spliced genes (Handa *et al*., 1996; Kim *et al*., 2016; Choi *et al*., 2020). The breakpoints of *ccmFn* differ between the three lineages, implying independent gene breakup events. In Convolvulaceae, the intronic *ccmFc* has encountered an unusual gene fission and *trans*-splicing shift, resulting in a rare three-partite structure in three different places on the mitochondrial chromosome (Lin et al., 2022).

In hornworts, lycophytes, and ferns—the latter including leptosporangiates—the loss of mitochondrial-encoded *ccm* genes must have occurred independently, which is suggested by their presence in the eusporangiate *Psilotum* (Guo *et al*., 2017)12/23/22 12:36:00 PM. The losses of *ccm* genes need not necessarily be the result of functional gene transfer from the mitochondrial to the nuclear genome. Although another eusporangiate fern, *Ophioglossum*, lacks mitochondrial *ccm* genes, no nuclear-transcribed copies could be identified (Guo *et al*., 2017). *Selaginella*, a *ccm*-lacking lycophyte species also encodes no nuclear-encoded *ccm* homologues (Guo *et al*., 2016). Hence, the entire *ccm* pathway might have been lost from these species, perhaps in favor of an alternative cytochrome maturation pathway (Guo *et al*., 2017).

#### Genes for other mitochondrial processes

Besides OXPHOS-related proteins, plant mitogenomes encode elements for translation, including three ribosomal (*rrn*) and several transfer RNAs (tRNAs), as well as various proteins for the large and small ribosomal subunit (*rpl* and *rps* genes, respectively). The plant mitogene set is complemented by open reading frames for proteins involved in RNA and protein processing.

The TatC protein functions in the twin-arginine translocation pathway for import of fully folded proteins (Carrie *et al*., 2016). *MttB* (transport membrane protein B, also named *orfX* or *ymf16*) is intact in all land plant mitogenomes (Figure 3; Table S1). In *Ipomoea* and *Cuscuta*, the *mttB* genes has a premature stop codon, which is not corrected by RNA editing (Figure S4), but perhaps through use of an alternative start codon. *MttB* is also speculated to be a pseudogene in some *Viscum* species (Petersen *et al*., 2015).

Encoding a pre-RNA maturase for splicing of mitochondrial group II introns (Sultan *et al*., 2016), the *matR* gene is specific to land plant mitogenomes. Nearly full-length copies exist in most mosses (Dombrovska and Qiu, 2004), hornworts (Guo and Mower, 2013), the lycophyte *Huperzia* (Liu *et al*., 2012), and the eusporangiate fern *Botrychium* (Davis *et al*., 2005). The leptosporangiate fern *Pteridium* retains *matR*, too, but it shows multiple premature stop codons and it is the only gene for which we detected no transcripts. Plants lacking mitochondrial *MatR* might use another maturase. Several mitochondrial ORFs similar to RNA maturases of yeast were detected in the liverwort *Marchantia* (Oda *et al*., 1992) and the lycophyte *Isoëtes* (Grewe *et al*., 2009). However, it remains to be experimentally validated whether the enigmatic pattern of *matR* gene losses in non-seed plants can be attributed to an alternative RNA maturase. In seed plants, *matR* is normally intact (Figure 3 and Supporting Table S1), although localized losses are known in *Viscum* (Skippington *et al*., 2015)12/23/22 12:36:00 PM, *Pelargonium* (Grewe *et al*., 2016), *Croizatia* and *Lachnostylis* (both Phyllanthaceae) (Wurdack and Davis, 2009). There are also several notable rearrangements around the *matR* gene, including in *Cuscuta* (Figure S4).

#### Genes for mitochondrial protein biosynthesis

Mitochondrial ribosomal proteins of plants are substantially divergent from those of bacteria and other eukaryotes. Land plant mitogenomes encode up to five proteins for the large ribosomal subunit (*rpl* genes *2, 5, 6, 10*, and *16*) and 12 proteins for the small one (*rps* genes *1*–*4, 7, 8, 10*–*14, 19*). Liverwort mitogenomes contain all of these 17 ribosomal proteins, whereas hornworts retain only two (*rpl10* and *rps13*). The *rps8* was only found in liverworts, and *rpl6* has been lost in the common ancestor of seed plants (Figure 3 and Supporting Table S1). Within lycophytes, the spikemoss *Selaginella* has lost all mitochondrial *rpl* and *rps* genes (Hecht *et al*., 2011), similar to *Isoëtes* that retains four (Grewe *et al*., 2009). *Huperzia*, ferns, gymnosperms (except for *Welwitschia*: Guo *et al*., 2016), and basal angiosperms contain an average of seven *rpl*/*rps* genes (Figure 3 and Supporting Table S1). In other angiosperms, the *rp*-gene content is very dynamic (Adams *et al*., 2002) (Table 1). Some species like *Zostera* (Petersen *et al*., 2017), *Allium* (Kim *et al*., 2016), *Geranium* (Park *et al*., 2015), *Viscum* (Petersen *et al*., 2015; Skippington *et al*., 2017; Skippington *et al*., 2015), *Silene* (Sloan, Alverson, Chuckalovcak, *et al*., 2012; Sloan, Alverson, Wu, *et al*., 2012; Štorchová *et al*., 2018), and *Ajuga* (Zhu *et al*., 2014) encode almost no ribosomal proteins in their mitogenomes, all of which retaining only the *rpl5, rps3, rps4*, and *rps12* genes. The loss of *rps2* and *rps11* might be a synapomorphy of most eudicots, where these are present only in *Nelumbo* (Nelumbonaceae: Gui *et al*., 2016). The *rps11* and *rps2* in *Betula* and *Actinidia*, respectively, have been acquired by horizontal gene transfer (HGT) events (Bergthorsson *et al*., 2003).

#### Structural RNAs

Three ribosomal RNAs—5S, 18S, and 26S—are encoded by the mitochondrial *rrn5*, rrn18 (*rrnS*), and *rrn26* (*rrnL*), respectively. Unlike ribosomal protein genes, rRNA genes are highly conserved in plants (Supporting Table S1). Mistletoes (*Viscum* sp.) might be the only lineage of land plants where *rrn* genes have been lost (Skippington *et al*., 2015; Petersen *et al*., 2015; Skippington *et al*., 2017).

Land plant mitochondria contain no complete set of tRNA isoacceptors for the 21 standard amino acids. That is, amino acids encoded by different codons still have only one native mitochondrial tRNA isoacceptor per amino acid–if at all (Supporting Table S1). Only marchantoid liverworts seem to retain a full mitochondrial set of 21 tRNA genes. Especially in vascular plants, tRNA gene losses are common. *Selaginella* apparently encodes not a single tRNA in its mitogenomes (Hecht *et al*., 2011), although other lycophytes, as well as ferns and seed plants, typically contain a least a dozen different transfer RNA genes (Table S1). Significant independent reductions of tRNA gene content has occurred in *Silene* (Sloan, Alverson, Chuckalovcak, *et al*., 2012) and *Viscum* (Skippington *et al*., 2015; Petersen *et al*., 2015; Skippington *et al*., 2017). Isoacceptors for tRNAs not encoded in plant mitochondrial genomes are likely imported from the cytosol (Dietrich *et al*., 1996; Glover *et al*., 2001). Additionally, mitochondrial DNA of plastid origin contain plastid-like tRNA genes of functional relevance for mitochondrial translation, even though the majority of plastid inserts into mitogenomes originate from otherwise nonfunctional intracellular DNA transfers (Small, 1999; Sloan *et al*., 2010).

### RNA editing

RNA editing—the conversion of cytidines to uridines (C-to-U) and uridines to cytidines (U-to-C) in transcripts of plant organelles (Castandet and Araya, 2011; Schallenberg-Rüdinger and Knoop, 2016)—often rescues putative pseudogenes and restores the gene product’s functionality. RNA editing may reinstate a proper start or stop codon, override premature stop codons, and affect RNA processing, and has the potential to generate new plant phenotypes (Small *et al*., 2019 for details).

The extent of RNA editing is highly lineage-specific. In spite of many RNA edit site predictions, a lack of actual cDNA or RNA evidence hampers robust conclusions. Liverworts and mosses post-transcriptionally edit only a relative small numbers of sites (up to a hundred) and fewer than predicted (Hiesel *et al*., 1994; Liu *et al*., 2011; Rüdinger *et al*., 2012). Marchantiid liverworts have even secondarily lost organellar RNA editing (Groth-Malonek *et al*., 2007; Rüdinger *et al*., 2012). The gymnosperms *Cycas* and *Ginkgo* each count more than 1,000 predicted edit site, five times more than *Welwitschia* (Chaw *et al*., 2008; Guo *et al*., 2016; Fan *et al*., 2019), which lies within the angiosperm range of 200 to 700 RNA edit sites (Edera *et al*., 2018).

Extreme high level of RNA editing occurs in hornworts, lycophytes, and ferns. Many mitochondrial genes require the conversion of multiple internal stop codons to normal codons. With over 2,400 edited sites, the current record holders of RNA corrections is the hornwort *Anthoceros* (followed by the lycophytes *Isoëtes* and *Selaginella* with ∼1,700 and 2,100 editing sites, respectively – Hecht *et al*., 2011; Grewe *et al*., 2011). With ∼800 RNA edit sites, the mitogenomes of two eusporangiate ferns (Guo *et al*., 2017) have slightly fewer edit site than leptosporangiate *Azolla* and *Pteridium*, for which we counted ∼1,300 and 1,600 edits, respectively, by RNA-DNA comparisons (Supporting Table S2).

Besides a canonical C-to-U editing, the reverse U-to-C editing is particularly frequent in hornworts (Villarreal *et al*., 2018; Gerke *et al*., 2020), lycophytes (Hecht *et al*., 2011; Hecht *et al*., 2011; Grewe *et al*., 2011), and ferns (Varigerow *et al*., 1999; Bégu *et al*., 2011; Guo *et al*., 2017; this study), where it represents about 20 % of the total RNA editing events. Reverse RNA editing is absent from mosses and gymnosperms, and it is extremely rare in liverworts and angiosperms (Gualberto *et al*., 1990; Schuster *et al*., 1990; Picardi *et al*., 2010). Our own analysis of *Azolla* and *Pteridium* indicates that U-to-C editing may reach its highest overall level in leptosporangiate ferns with a percentage of 30 and 40%, respectively—the second highest in land plants after hornworts (∼54 %, Gerke *et al*., 2020). These results corroborate that C-to-U and U-to-C editing might (have) evolve(d) independently of and disproportionally to one another in these two lineages (Knie *et al*., 2016), and/or that these plants contain a unique RNA editing machinery.

### Molecular evolution

Despite their dynamic structural evolution, mitochondrial genomes evolve slower than all other cellular genomes in plants (Palmer and Herbon, 1988). The majority of genes evolve at an overall low rate of molecular evolution. Even across land plants, elevations of substitution rates are limited to either single genes, and indel events in coding regions are rare (Supporting Figure S4). Across the sampled 218 land plants (i.e., excluding the charophyte algae), we detected a significant difference in substitution rates between many *nad, cox* and *cob* genes and others (non-parametric Wilcoxon Rank Sum test, P-value <0.001) corroborating earlier findings of gene class-specific nonsynonymous and synonymous rates of evolution in organellar genomes (Wicke and Schneeweiss, 2015). Despite an overall low nonsynonymous substitution rate, we observe several accelerations and decelerations in a few genes like *ccmFc, nad5, nad6, nad9, matR*, or *rps14* (Figure 3). In contrast, synonymous rates seem to experience fewer ups and downs during the evolution of land plants, although genes like *cox3, rps3*, and *mttB* show more synonymous changes than most other mitochondrial genes. Within angiosperms, we observe no notable differences of substitution rates between asterids and rosids, even though their structural mitogenomic traits diverge marginally from one another (Figure 1 and 2).

Non-seed plants show a slightly though significantly higher rate of synonymous substitutions across all gene classes (Figure 3). In contrast, vascular plants appear to evolve under slightly higher non-synonymous rates in all genes than bryophytes. Despite a considerable variation of evolutionary rates within the various land plant lineage (Figures 3 and 4), we observe significant shifts of both dN and dS between bryophytes and vascular plants but also between non-seed and seed plants (all P-values <0.001;). Having the broadest range of substitution rates (Figure 4), ferns differ significantly in the number of nonsynonymous and synonymous changes compared with all non-seed plant and angiosperms in their molecular evolution. This pattern of substitution rate evolution is not unexpected, as it matches the outstandingly high rates of RNA editing-based nucleotide corrections (this study; Knie et al. 2016). The similarly high editing rates in lycophytes are also reflected in a broad substitution rate range, but a significant difference in molecular evolutionary rate between lycophytes and bryophytes cannot be detected (Figure 4). Our analyses suggest that the change of molecular evolutionary rates (dS) are most notably positively associated with shifts of nucleotide composition (posterior probability [pp]: 0.95; corrected for co-correlations based on 30,000 effective samples) and negatively correlate with mitogenome size (pp: 0.01). The latter itself relates strongly to repeat length (pp: 1.00). Marginal correlations with the percentage of protein-coding regions and structural RNAs cannot hold corrections for co-correlations (Table 2). Together, these results reflect the considerable changes of mito-chromosomal architecture and might be related to increases of error-prone repair mechanisms.

**Figure 4.**
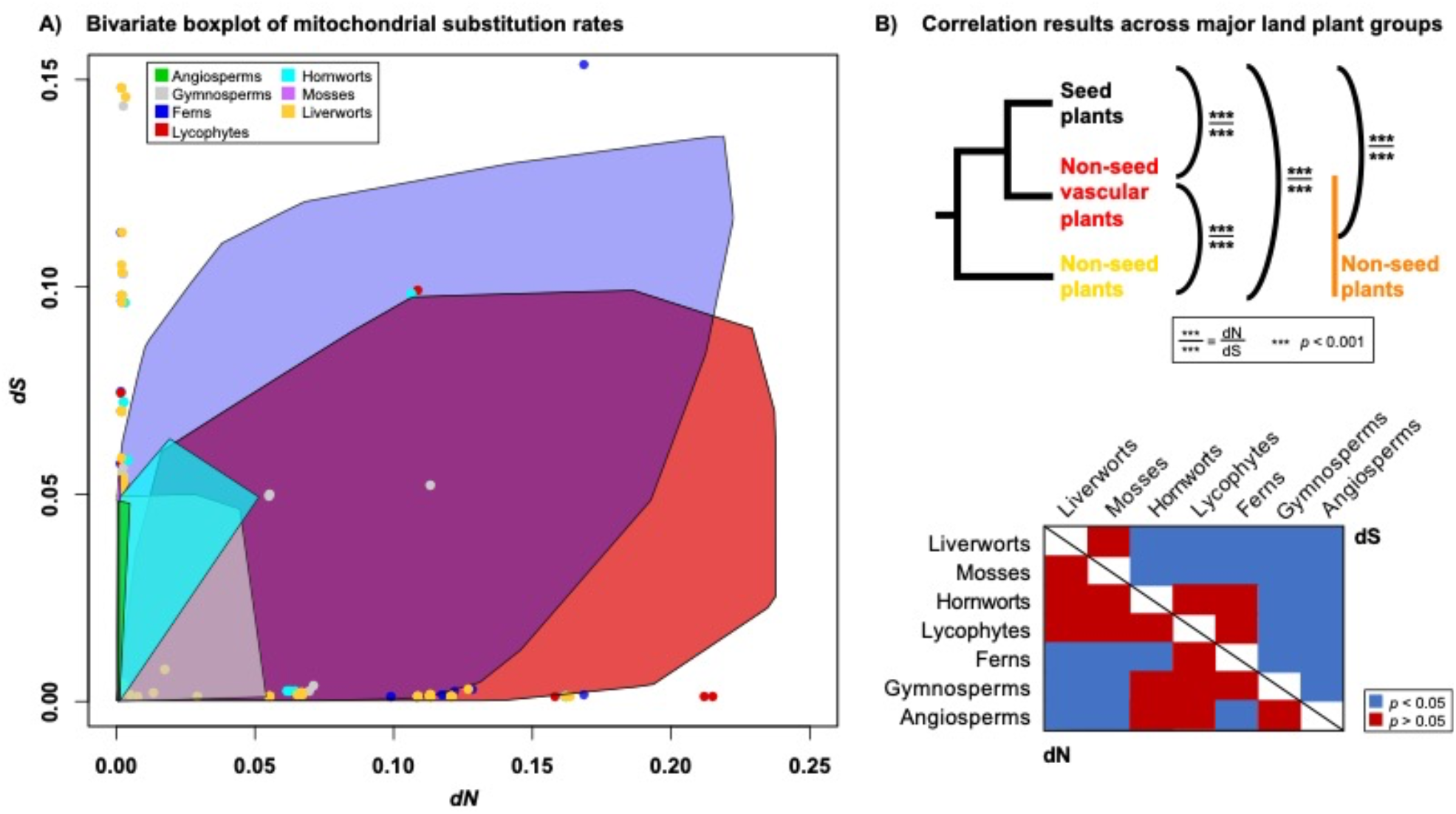
Comparison of mitochondrial nucleotide substitution rates of the major groups in land plants. **A**. Boxplot bag hulls illustrate the range of relative substitution rates (dN, dS) across all mitogenes per land plant lineage, the identity of which is color-coded. Bag hulls indicate the 25^th^ and 75^th^ percentiles of the upper and lower bounds of the lineage-specific range of substitution rates. **B**. The differences of molecular evolutionary rates between the various land plant lineages is graphically summarized on the tree, with three asterisks indicating a p-level < 0.001 of statistical significance, whereby dN is shown above and dS below branches. Lineage-wise comparisons are graphically summarized in tabulated format, highlighting significant differences in blue, whereas red indicates no differences in dS (upper-right triangle) or dN (lower-left triangle), respectively.

Being known for their extraordinarily elevated substitution rates in plastid genomes (Wicke *et al*., 2016; Wicke and Naumann, 2018), parasitic plant mitogenomes show apparently no lifestyle-related regime of molecular evolution (Figure 3). Although some parasitic angiosperms within asterids undergo individual routes of genomic evolution, none of the parasitic species stand out by long branches (Figure 1) or a higher number of genes with elevated non-synonymous or synonymous rates (Zervas *et al*., 2019). It is likely that the transition from an autotrophic to a heterotrophic way of life, where mitochondrial energy production is less affected by the altered lifestyle, might have only marginal effect on mitogenomic evolution. Thus, observations that “parasitic plants have increased rates of molecular evolution across all three genomes” (Bromham *et al*., 2013, p.126) might not hold up under rigorous hypothesis testing involving the entire gene sets. However, parasitism has been associated with higher rates of horizontal gene transfer in mitochondrial genomes as well as with elevated rates in a few genes (Nickrent and Starr, 1994; Duff and Nickrent, 1997; Barkman *et al*., 2007; Bromham *et al*., 2013; Xi *et al*., 2013), compared with their respective closest relatives. Future studies considering different forms of trophic specializations are needed to clarify the extent of parasitism-related changes of mitogenome evolution.

## Conclusion

Here, we presented an analysis of mitogenome evolution across land plants, thereby providing a thorough review of existing knowledge alongside a novel, a priori-free evaluation of molecular evolutionary and genetic traits based on an unprecedent data matrix. In doing so, this work also provides curated land plant-wide data sets of all mitochondrial genes for 218 taxa (https://github.com/wickeLab) as we all as a compilation of mitogenomic traits for over 300 taxa (Supporting Table S1). This work also fills the gap of lacking leptosporangiate fern mitogenomes by reconstructing the mitochondrial genomes of *Azolla* and *Pteridium*, including their RNA editing behavior.

In sum, this work highlights the extraordinarily diverse evolutionary trajectories of mitochondrial chromosomes in land plants. The small and compact bryophyte mitogenomes still resemble their green algal ancestors. The evolution of vascularity brings about notable changes in mitogenomic architecture and coding capacity as well as coinciding shifts of molecular evolution, which in turn reach their maxima in lineages with the emergence of (extreme) RNA editing. Although also lycophytes exhibit dramatic levels of RNA editing, fern mitogenomes seem to depart from all other land plants by reaching more extreme levels of both C-to-U and U-to-C editing. As a result, genic conservation decreases in ferns, resulting in highly diverged coding regions, which potentially have hampered past efforts to sequence the mitochondrial genome using homology-based approaches. Unravelling the mechanisms of the latter in ferns in lycophytes and how that differs from the genetic makeup of other land plants might contribute new paths for a targeted, post-transcriptional manipulation of plant functions, e.g., crops. These molecular-evolutionary departure with extreme sequence drift might have led to leptosporangiate ferns being the last major plant lineages for which mitogenomic evidence becomes available. The ferns’ extant sister group—seed plants—also shows extreme departures from the canonical mitogenomic evolution of early-diverging land plants. Especially in angiosperms, mitogenomes physically inflate multiple times independently (Figures 1 and 2) despite several functional reductions (*rpl, rps*, tRNAs, *sdh* genes) and the mito-chromosomal architectures often exhibit an enormous complexity of DNA mosaics.

This study and the associated data collection might be helpful for future research of mitochondrial and their genetic material. As energy factory of a cell, knowledge of their plant lineage-specific peculiarities is valuable for in-depth functional evaluations, to the extent of rescuing CMS in crops. Unlike plastid genomes but similar to the more untractable nuclear genomes, the dynamically evolving mitogenomes of plants knowingly retain the ability to generate new genes or isoforms. It will be worthy of further study to investigate in high-throughput assays the transcriptional and translationally activities in mitogenomes of various land plant lineages, especially of non-seed plants. Such research might lead to detecting newly evolved genes or highly conserved regulatory elements. The highly dynamic and diverse mitogenomes in land plants also offer an excellent model genome to understand mechanism of genomic change. Compared with many plant nuclear genomes, there are more compact and present in high copy numbers and experience recombination. Studied in an explicit phylogenetic framework, mitogenomes can contribute to understanding the mechanisms of rearrangements, physical DNA losses and gains, intronization, repeat proliferation, and provide new clues about organellar double strain breaks and repairs. To conclude, further study of mitochondria and their genetic repertoire in plants is essential, and more data covering all land plant lineages and from more than one individual per group is needed.

## Data availability

The newly assembled and annotated genomes have been deposited at NBCI Genbank under the following accession numbers: MN400566-MN400574 (*Azolla*), and MN400594–MN400606 (*Pteridium*), MN400575 MN400593 (*Cuscuta* spp.). Other information and scripts used herein are available at Mendeley Data, doi: 10.17632/4fgcswrw3v.1.

## Acknowledgments

This study received support from CAS in the form of a doctoral scholarship to Y.F. and the German Science Foundation to S.W. (DFG, WI4507/9-1).

