## Supplementary material for "New mitochondrial genomes of leptosporangiate ferns allow modeling the mitogenomic inflation syndrome across all land plant lineages": Table S2

**Table S2. RNA editing sites in land plants, showing U-to-C as main numbers and U-to-C edits in round brackets.**

| | Lei | Sel | Iso | Oph | Psi | Azo | PSC | Pte | Gin | Wel | Lir | Ory | Ara | Nic | Ipo | Cau | Cca |
|---|---|---|---|---|---|---|---|---|---|---|---|---|---|---|---|---|---|
| *atp1* | 2 | 160 | 131 (27) | 8 | 10 | 99 (51) | 29 [6] | 92 (43) | 23 | | 14 | 5 | 5 | 7 | 2 | 2 | 2 |
| *atp4* | 1 | | 6 | 17 | 10 | 33 (17) | 13 | 38 (15) | 27 | | 16 | 9 | 8 | 10 | 9 | 8 | 8 |
| *atp6* | 12 (1) | 80 | 95 (12) | 49 (1) | 26 | 62 (22) | 8 [3] | 51 (16) | 63 | | 29 | 17 | 1 | 19 | 22 | 20 | 20 |
| *atp8* | | 35 | 35 (4) | 9 (1) | 9 | 20 (5) | 8 [3] | 23 (8) | 19 | | 6 | 4 | | 6 | 5 | 6 | 6 |
| *atp9* | 2 | 44 | 34 (4) | 16 (1) | 14 | 20 (7) | 2 [2] | 19 (4) | 21 | | 15 | 8 | 4 | 11 | 9 | 8 | 8 |
| *ccmB* | | | | | 20 (1) | | | | 21 | 19 | 49 | 35 | 39 | 48 | 23 | 18 | 28 |
| *ccmC* | | | | | 35 | | | | 45 | 3 | 41 | 36 | 28 | 37 | 31 | 31 | 28 |
| *ccmFc* | | | | | 31 | | | | 14 | 11 | 24 | 27 | 16 | 21 | 12 | 10 | 16 |
| *ccmFn* | | | | | 36 (2) | | | | 35 | | 34 | 31 | 34 | 34 | 9 | 17 | 14 |
| *cob* | 8 (1) | 122 | 121 (22) | 42 (6) | 17 (1) | 93 (43) | 11 [2] | 88 (39) | 3 | | 30 | 19 | 7 | 14 | 8 | 9 | 9 |
| *cox1* | 2 | 181 | 110 (22) | 32 | 41 (1) | 129 (45) | 11 [5] | 119 (44) | 3 | | 37 | 4 | | 18 | 16 | 17 | 17 |
| *cox2* | | 97 | 14 (4) | 14 | 17 | 46 (24) | 12 [2] | 49 (22) | 27 | | 14 | 19 | 15 | 15 | 15 | 14 | 14 |
| *cox3* | 3 (2) | 133 | 101 (3) | 28 (1) | 18 | 40 (16) | 5 | 44 (13) | 46 | | 19 | 1 | 8 | 9 | 11 | 11 | 11 |
| *matR* | | | 77 (5) | 1 | 1 | | | 0 | 31 | 14 | 19 | | 9 | 15 | 10 | 13 | 14 |
| *mttB* | 10 | 133 | 68 (6) | 29 | 40 | 56 (14) | 5 [2] | 7 (3) | 37 | 10 | 58 | 33 | 24 | | 10 | 26 | 28 |
| *nad1* | 7 (4) | 137 | 57 (9) | 41 (5) | 28 (2) | 75 (29) | 8 [1] | 66 (25) | 38 | 2 | 33 | 23 | 24 | 28 | 13 | 19 | 20 |
| *nad2* | 10 (1) | 249 | 100 (6) | 66 (4) | 29 (1) | 75 (20) | 6 | 70 (15) | 57 | | 49 | 30 | 32 | 40 | 17 | 26 | 26 |
| *nad3* | 4 | 77 | 57 (4) | 23 (2) | 20 | 33 (7) | 1 [1] | 20 (2) | 7 | | 21 | 15 | 12 | 18 | 16 | 12 | 12 |
| *nad4* | 3 | 237 | 156 (16) | 82 (5) | 56 (1) | 92 (38) | 14 [2] | 92 (20) | 103 | 1 | 58 | 20 | 32 | 44 | 34 | 36 | 36 |
| *nad4L* | 4 (2) | 47 | 27 (1) | 21 (2) | 10 | 25 (5) | 1 | 23 (4) | 20 | | 13 | 10 | 9 | 16 | 13 | 14 | 14 |
| *nad5* | 13 (4) | 143 | 166 (14) | 57 (5) | 43 (1) | 107 (38) | 11 [1] | 70 (16) | 114 | 2 | 49 | 11 | 27 | 29 | 23 | 21 | 21 |
| *nad6* | 5 | 89 | 64 (6) | 29 (1) | 14 | 26 (4) | 1 | 25 (1) | 47 | 22 | 25 | 18 | 10 | 7 | 7 | 8 | 7 |
| *nad7* | 1 (1) | 103 | 115 (20) | 40 (7) | 35 (2) | 91 (43) | 27 | 79 (32) | 45 | 9 | 39 | 32 | 27 | 30 | 23 | 26 | 26 |
| *nad9* | | 72 | 50 (10) | 22 (5) | 14 | 50 (22) | 6 | | 25 | | 14 | 12 | 7 | 8 | 9 | 7 | 7 |
| *rpl2* | | | | 12 (1) | 9 | 49 (34) | 18 | 33 (19) | 13 | | 3 | 1 | 1 | | | | |
| *rpl5* | | | 30 (7) | 6 (1) | 8 (1) | 34 (19) | 10 | 26 (11) | 17 | | 12 | 1 | 10 | | 9 | 7 | 8 |
| *rpl6* | | | | 11 | 7 | 18 (8) | 8 [2] | 20 (9) | | | | | | | | | |
| *rpl10* | | | | | | | | | 12 | 1 | | | | | 7 | | |
| *rpl16* | | | | 10 (1) | 8 | 18 (11) | 6 | 7 (6) | 11 | | | 12 | 5 | | 5 | 2 | 2 |
| *rps1* | | | | 17 | 10 | 21 (7) | 1 [1] | 13 (3) | 27 | | | 3 | | | 3 | 3 | 3 |
| *rps2* | | | 28 (5) | 10 (1) | 11 | 31 (22) | 9 | 25 (7) | 9 | | 17 | 10 | | | | | |
| *rps3* | | | 62 (6) | 28 | 17 (2) | 59 (31) | 20 | 46 (13) | 33 | 1 | | 10 | 13 | | 11 | 8 | 7 |
| *rps4* | | | 55 (7) | 23 | 24 | 42 (17) | 13 [1] | 20 (11) | 47 | 4 | | 15 | 15 | | 8 | | |
| *rps7* | | | | 15 | 6 | 15 (7) | 5 | 16 (8) | 18 | | 4 | 2 | | | | | |
| *rps10* | | | | 4 | | 21 (11) | 6 | 18 (10) | 2 | | 6 | | | | 2 | | |
| *rps11* | | | | 23 (2) | 12 | 39 (25) | 12 | 29 (17) | 19 | | | 4 | | | | | |
| *rps12* | | | | 11 | 9 | 34 (15) | 9 [2] | 18 (9) | 18 | | 9 | | 8 | 4 | 7 | 5 | 5 |
| *rps13* | | | | 8 | 7 | 23 (14) | 10 | 28 (15) | 2 | | 7 | 8 | | | 4 | 5 | 9 |
| *rps14* | | | | 12 (3) | 6 (1) | 12 (8) | 4 [2] | 9 (3) | 11 | | 2 | | | | 1 | 0 | 0 |
| *rps19* | | | | 5 | 4 | 9 (6) | 3 | 8 (5) | 13 | | 5 | | | | 3 | 0 | 0 |
| *sdh3* | 5 (1) | | 23 (2) | 28 (1) | 11 | 21 (9) | 5 [1] | 11 (4) | 23 | | 10 | | | | | | |
| *sdh4* | 1 | | | 13 | 8 | 16 (4) | 2 | 12 (3) | 16 | | | | | | 8 | | |
| Total | 93 (17) | 2139 | 1782 (222) | 862 (56) | 731 (16) | 1634 (698) | 320 [39] | 1314 (475) | 1162 | 99 | 781 | 491 | 430 | 488 | 415 | 409 | 426 |

The column PSC after *Azolla* shows premature stop codons in *Azolla*, numbers in square brackets mean the stop codons that are likely not corrected by RNA editing. Lei: Leiosporoceros: (Villarreal et al., 2018), Sel: Selaginella: (Hecht et al., 2011), Iso: Isoetes: (Grewe et al., 2011), Oph: Ophioglossum and Psi: Psilotum: (Guo et al., 2017), Gin: Ginkgo and Wel: Welwitschia: (Fan et al., 2019), Lir: Liriodendron: (Richardson et al., 2013), Ory: Oryza: (Notsu et al., 2002), Ara: Arabidopsis: (Giegé and Brennicke, 1999), Nic: Nicotiana: (Edera et al., 2018).

Edera, A.A., Gandini, C.L., and Sanchez-Puerta, M.V. (2018). Towards a comprehensive picture of C-to-U RNA editing sites in angiosperm mitochondria. Plant Mol Biol *97*, 215-231.

Fan, W., Guo, W., Funk, L., Mower, J.P., and Zhu, A. (2019). Complete loss of RNA editing from the plastid genome and most highly expressed mitochondrial genes of Welwitschia mirabilis. Sci China Life Sci *62*, 498-506.

Giegé, P., and Brennicke, A. (1999). RNA editing in Arabidopsis mitochondria effects 441 C to U changes in ORFs. P Natl Acad Sci USA *96*, 15324-15329.

Grewe, F., Herres, S., Viehover, P., Polsakiewicz, M., Weisshaar, B., and Knoop, V. (2011). A unique transcriptome: 1782 positions of RNA editing alter 1406 codon identities in mitochondrial mRNAs of the lycophyte Isoetes engelmannii. Nucleic Acids Res *39*, 2890-2902.

Guo, W., Zhu, A., Fan, W., and Mower, J.P. (2017). Complete mitochondrial genomes from the ferns Ophioglossum californicum and Psilotum nudum are highly repetitive with the largest organellar introns. New Phytol *213*, 391-403.

Hecht, J., Grewe, F., and Knoop, V. (2011). Extreme RNA Editing in Coding Islands and Abundant Microsatellites in Repeat Sequences of Selaginella moellendorffii Mitochondria: The Root of Frequent Plant mtDNA Recombination in Early Tracheophytes. Genome Biol Evol *3*, 344-358.

Notsu, Y., Masood, S., Nishikawa, T., Kubo, N., Akiduki, G., Nakazono, M., Hirai, A., and Kadowaki, K. (2002). The complete sequence of the rice (Oryza sativa L.) mitochondrial genome: frequent DNA sequence acquisition and loss during the evolution of flowering plants. Mol Genet Genomics *268*, 434-445.

Richardson, A.O., Rice, D.W., Young, G.J., Alverson, A.J., and Palmer, J.D. (2013). The "fossilized" mitochondrial genome of Liriodendron tulipifera: ancestral gene content and order, ancestral editing sites, and extraordinarily low mutation rate. BMC Biol *11*, 29.

Villarreal, A.J., Turmel, M., Bourgouin-Couture, M., Laroche, J., Salazar Allen, N., Li, F.W., Cheng, S., Renzaglia, K., and Lemieux, C. (2018). Genome-wide organellar analyses from the hornwort Leiosporoceros dussii show low frequency of RNA editing. Plos One *13*, e0200491.
