## Supporting Figure for "New mitochondrial genomes of leptosporangiate ferns allow modeling the mitogenomic inflation syndrome across all land plant lineages"

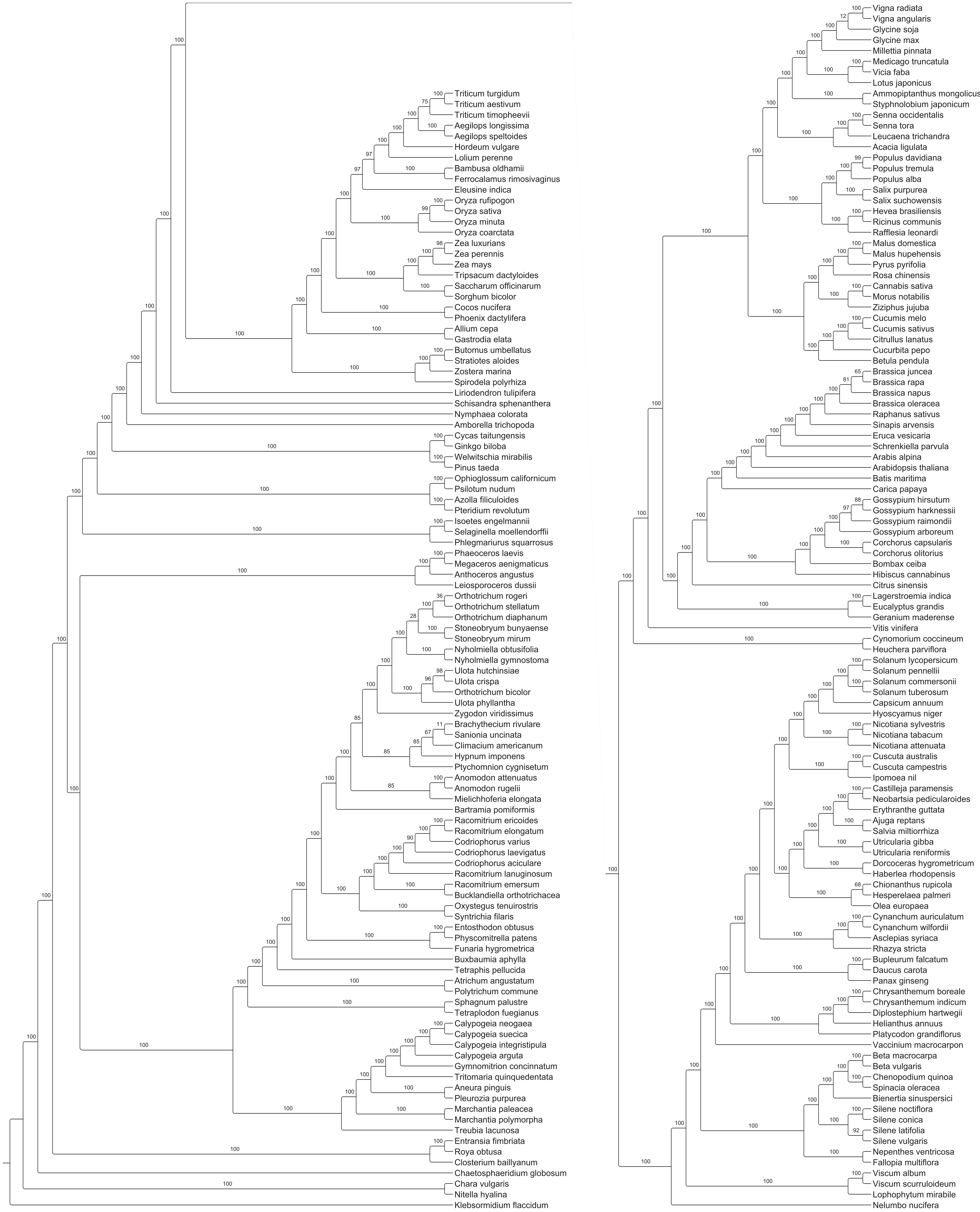


**Figure S1 Phylogenetic tree resulting from an analysis of land plant-wide mitochondrial genome data and maximum likelihood inferrence.** Support values above nodes result from 1,000 bootstrap iterations.

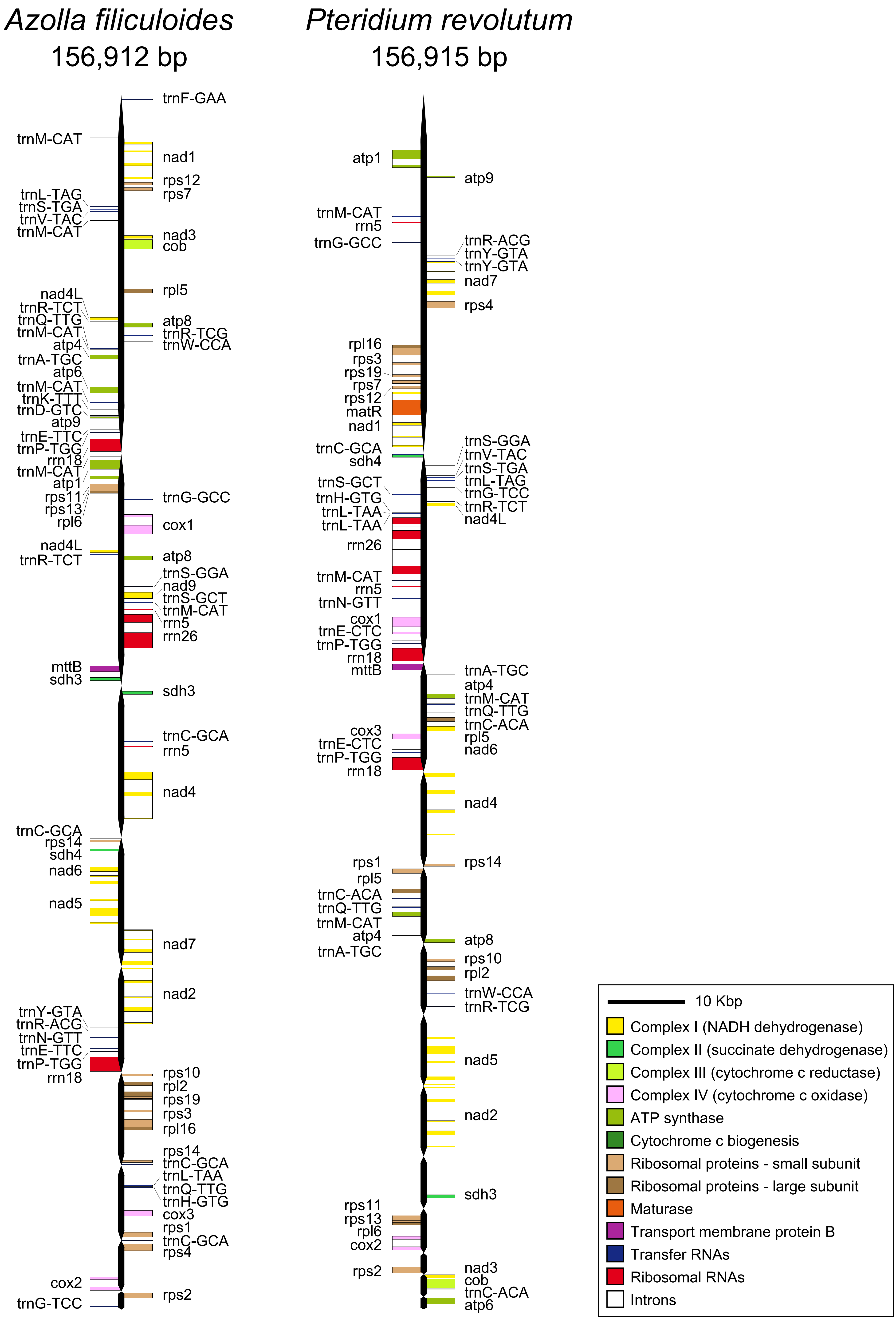


**Figure S2 Physical maps of the reconstructed mitochondrial genomes of *Azolla* and *Pteridium*.** Contiguous fragments with gene blocks are shown in black bars; contig breaks are indicated by a thinning of the chromosomal bar.

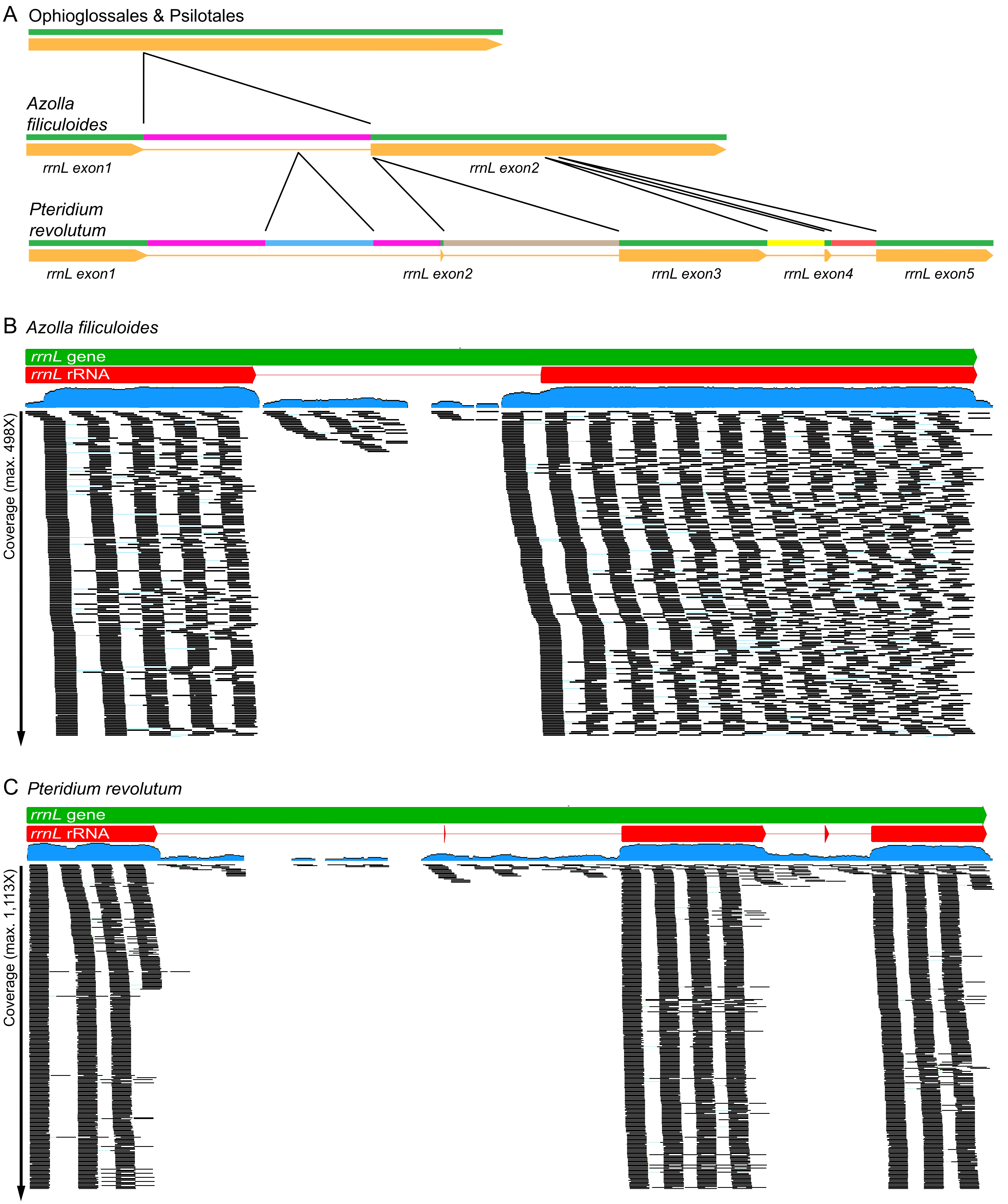


**Figure S3 Graphical summary of read mappings and divergence in the mitochondrial rrnL gene region. A.** Changes in intronization of the rrnL gene regions between two representatives of eusporangiate ferns, *Azolla*, and *Pteridium*. Read distribution across the mitochondrial rrnL gene region of **B.** *Azolla* and **C.** *Pteridium*. The rRNA coding region is indicated by red bars on top. The overall read mapping density is indicated in blue underneath the rRNA coding region. Blacks bars represent overlapping read stacks.

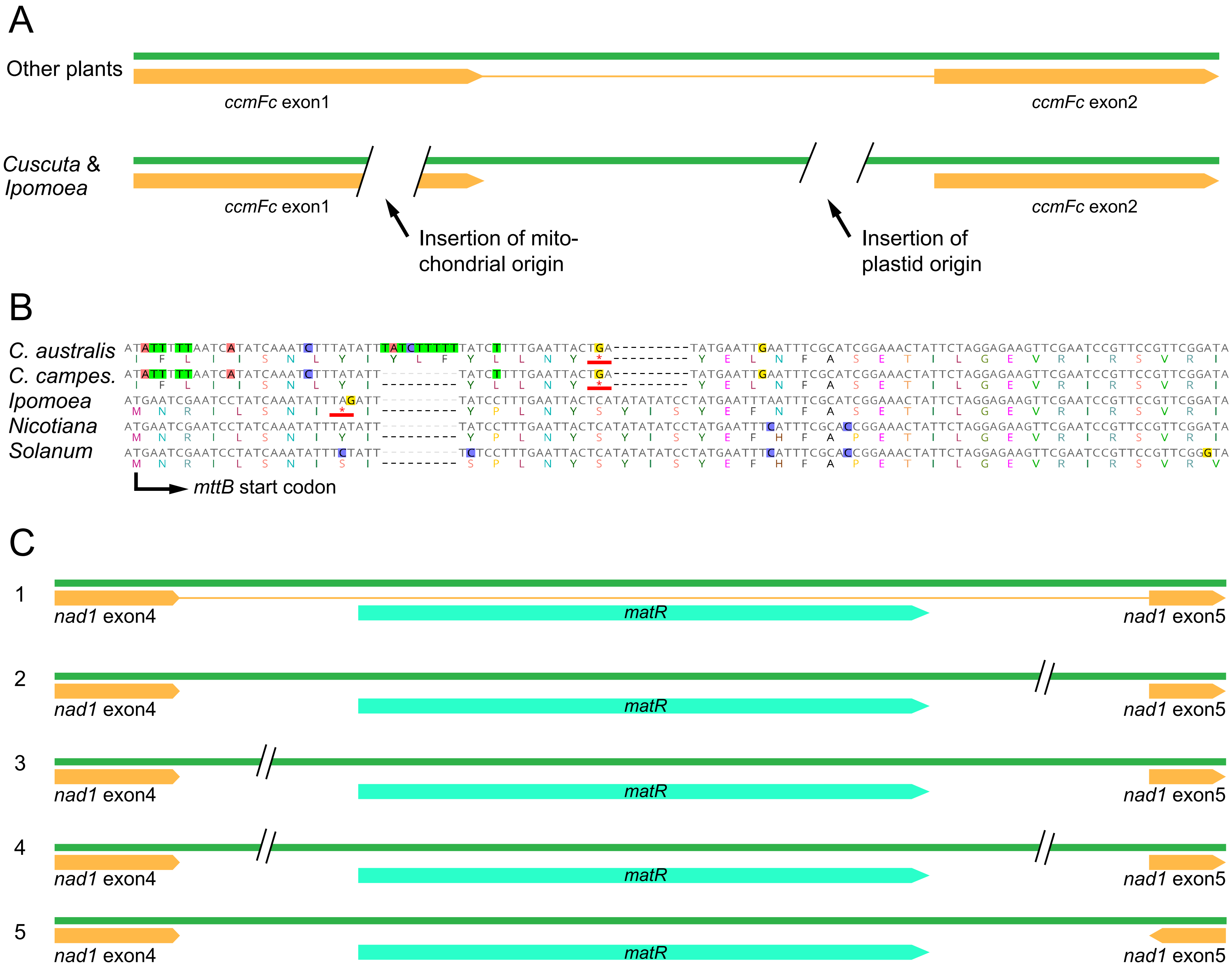


**Figure S4 Graphical summary of selected gene regions in Convolvulaceae. A.** Changes in the ccmFC coding region between Convulvulaceae and other plants. **B.** Evidence of putative pseudogenization within the *mttB* coding region of *Cuscuta* spp. and *Ipomoea* are internal stop codons (red underlined TGA or TAG codons, respectively. **C.** Reorganization of the nad1 coding region in Convolvulaceae, showing the nad1 exon 4 – matR – nad1 exon5 block variants. Case 1 represent the original state, which encountered different breaks in specific lineages as indicated by cases 2, 3, 4, and 5), the latter of which representing the condition in *Cuscuta* spp., where exon5 reside in reversed order.
